# Disrupted brain structural connectivity in Pediatric Bipolar Disorder with psychosis

**DOI:** 10.1101/241091

**Authors:** Henrique M. Fernandes, Joana Cabral, Tim J. Van Hartevelt, Louis-David Lord, Carsten Gleesborg, Arne Moller, Gustavo Deco, Peter C. Whybrow, Predrag Petrovic, Anthony C. James, Morten L. Kringelbach

## Abstract

Bipolar disorder (BD) has been linked to disrupted structural and functional connectivity between prefrontal networks and limbic brain regions. Studies of patients with pediatric bipolar disorder (PBD) can help elucidate the developmental origins of altered structural connectivity underlying BD and provide novel insights into the aetiology of BD. Here we compare the network properties of whole-brain structural connectomes of PBD patients with psychosis and euthymic matched healthy controls. Our results show widespread changes in the structural connectivity of PBD patients in both cortical and subcortical networks, notably affecting the orbitofrontal cortex, frontal gyrus, amygdala, hippocampus and basal ganglia. Graph theoretical analysis revealed that PBD connectomes have fewer hubs, weaker rich club organization, different modular fingerprint and inter-modular communication, compared to healthy participants. The relationship between network features and neurocognitive and psychotic scores was also assessed. Patients’ IQ and psychotic symptoms significantly correlated with the local efficiency of the orbitofrontal cortex. Our findings reveal that PBD is associated with significant widespread changes in structural network topology, thus strengthening the hypothesis of a reduced capacity for integrative processing of information across brain regions. Localised network changes involve core regions for emotional processing and regulation, as well as memory and executive function, some of which correlate with neurocognitive faculties and symptoms. Together, our findings provide the first comprehensive characterisation of the alterations in local and global structural brain connectivity and network topology, which may contribute to the deficits in cognition and emotion processing and regulation found in PBD.

## Introduction

Bipolar disorder (BD) is a psychiatric illness characterized by episodes of mania or hypomania and depression interleaved with euthymic periods. It has been suggested that prefrontal control over emotional networks is deficient in BD, inducing abnormally heightened emotional and reward processing in patients^1–4^.

Functional neuroimaging studies of euthymic bipolar patients have shown both reduced activity in the prefrontal cortex (PFC), especially in the ventrolateral and ventromedial PFC, and increased activity in emotional processing areas such as the ventral striatum^5^, the amygdala^6^ and the insula^7,8^ whilst performing a range of emotional processing tasks. Similar activity patterns in PFC were also observed in manic^3^ and depressed^9,10^ episodes. Moreover, functional connectivity studies have suggested a decreased coupling between frontal and limbic areas^10^, which might reflect abnormalities in the way these regions jointly process information in BD.

In contrast, structural neuroimaging studies are not dependent on the choice of an experimental task. The need for higher statistical power and convergence of the findings has motivated the production of several large voxel-based meta-analyses^11–14^, as well as large single-centre studies^15^. These structural studies have found decreased brain volume in BD in a specific set of brain regions including medial PFC, anterior cingulate cortex and ventrolateral PFC as well as the insula.

In addition to volumetric analyses, studies of structural connectivity (SC) in BD, which serves as the structural substrate over which the repertoire of functional networks can unfold, can provide highly valuable insights into the neural mechanisms underlying this disorder. Diffusion imaging studies have found altered between hemispheres^16,17^, as well as posterior cingulum^17^, prefrontal and subcortical regions in BD, which may result in abnormal communication between these regions and disrupt the modulation of emotion processing, onset of mania and the development of BD^2,18–20^.

In summary, there is emerging evidence linking BD to altered functional and structural connectivity between prefrontal regions and emotion processing regions such as the insula and the amygdala. BD most often starts in childhood or adolescence with sub-clinical or clinical manifestations of the disorder, suggesting an important developmental component^21^. Pediatric bipolar disorder (PBD) is a cyclic mood disorder in children and adolescents, which can be persistent and severe, and has an estimated prevalence of around 1%^22^. Few studies have explored the connectivity patterns in the neurodevelopmental phase of adolescence, when active brain reorganisation is occurring. Early-onset bipolar disorder with psychosis represents a useful, possibly, intermediate state to examine. Frontal connectivity alterations are common to both schizophrenia and bipolar disorder, with prominent fronto-temporal deficits identified in schizophrenia and inter-hemispheric and limbic alterations reported in bipolar disorder^23^.

Although abnormal functional connectivity is proposed to be a main feature in BD, few studies have focused on whether the underlying structural connectivity is altered in BD using state-of-the-art whole-brain network analysis^24–30^. As such, complementing the existing knowledge of anatomical and functional deficits in PBD with a comprehensive mapping of altered structural connectivity will inform how brain structure-function coupling is affected, and thus allow for an improved assessment of patients’ cognition^27^. Furthermore, this would help to minimise potentially confounding secondary manifestations of BD, which is key to elucidate which neural mechanisms are inducing and maintaining manic and depressed states in BD.

Here we constructed the structural connectomes of a group of adolescents with PBD and psychosis and of closely matched healthy controls. We used advanced structural connectomics analysis to characterise the topological differences in structural connectivity driving the cognitive and emotional symptoms found in PBD.

## Methods

### Participants

We analysed data from 15 patients with PBD with psychosis from the Oxford regional unit and surrounding units, and 15 euthymic age- and gender-matched healthy controls (HC) (Table 1). The patients were diagnosed according to the DSM-IV-TR criteria using the Kiddie Schedule for Affective Disorders and Schizophrenia (K-SADS-PL), and were administered the Positive and Negative Syndrome Scale (PANSS). The healthy subjects were recruited from the community through their general practitioners and interviewed using the K-SADS-PL to rule out any history of emotional, behavioural, or medical problems. Handedness was assessed with the Edinburgh Handedness Questionnaire. Intellectual ability was assessed using the Wechsler Abbreviated Scale of Intelligence (WASI). All subjects were clinically reviewed after the initial diagnostic screening for a period of at least six months (mean±std; 10±1.5 months); no subject changed diagnostic status. James and colleagues previously reported a comprehensive description of the cohort and clinical assessment strategy used in this study^2^. This study was undertaken in accordance with the guidance of the Oxford Psychiatric Research Ethics Committee (OPREC). All clinical and neuroimaging protocols included in this study were approved by the OPREC. Written consent was obtained from all participants and parents if subjects were under the age of 16.

**Table 1.** Demographic and clinical characteristics of participants.

| Characteristic | Normal Controls |  | Patients with PBD |  | p-value |
| --- | --- | --- | --- | --- | --- |
|  | mean | SD | mean | SD |  |
| Gender, M/F | 8/7 | - | 8/7 | - | 1 |
| Handedness, R/L | 13/2 |  | 13/2 |  | 1 |
| Age at onset of symptoms, years | 15.70 | 1.26 | 15.04 | 2.04 | 0.30 |
| Disease duration, years |  |  | 14.00 | 2.00 |  |
| Verbal IQ | 103.60 | 19.66 | 94.80 | 17.03 | 0.20 |
| Performance IQ | 106.53 | 12.98 | 99.13 | 15.01 | 0.16 |
| Full Scale IQ | 105.47 | 16.37 | 96.33 | 16.04 | 0.13 |
| Coding | - | - | 81.33 | 19.77 | - |
| PANSS: positive scores | - | - | 19.27 | 3.92 | - |
| PANNS: negative scores | - | - | 10.13 | 2.72 | - |
| Beck Depression Inventory |  |  | 6.60 | 1.40 |  |
| Young Mania Rating Scale |  |  | 1.40 | 0.80 |  |
| Comorbidity (ADHD), n |  |  | 5/15 |  |  |
| Medication, n |  |  |  |  |  |
| Olanzapine |  |  | 7 |  |  |
| Quetiapine |  |  | 3 |  |  |
| Risperidone |  |  | 1 |  |  |
| Fluoxetine |  |  | 1 |  |  |
| Sodium valproate |  |  | 2 |  |  |
| Lithium |  |  | 3 |  |  |

PANSS = Positive and Negative Syndrome Scale; ADHD = attention-deficit hyperactivity disorder.

### Image acquisition

All 30 participants underwent the acquisition of whole-brain T1-weighted and diffusion-weighted images using a 1.5 T Sonata magnetic resonance imager (Siemens, Erlangen, Germany) with a standard quadrature head coil and maximum 40 mT ⁄ m gradient capability. The 3D T1-weighted FLASH sequence was performed with the following parameters: coronal orientation; 256×256 reconstructed matrix; 208 slices; 1×1 mm^2^ in-plane resolution; slice thickness of 1 mm; echo time (TE) of 5.6 ms; repetition time (TR) of 12 ms; flip-angle (α) of 19°. The diffusion-weighted sequences were obtained using echo-planar imaging (SE-EPI), and its scanning parameters were: TE of 89 ms; TR of 8500 ms; 60 axial slices; bandwidth = 1860 Hz ⁄vx; voxel size of 2.5×2.5×2.5 mm^3^); 60 isotropically distributed orientations for the diffusion-sensitising gradients at a b-value of 1000 s⁄mm^2^ and five b0 images. To increase signal-to-noise ratio, scanning was repeated three times and all scans were merged.

### Network construction

The construction of the brain structural network for each experimental group consisted of a two-step process. First, the nodes of the network were defined using a brain parcellation. Secondly, the connections between nodes (i.e. edges) were estimated using probabilistic tractography. In the following we outline the details involved in each step.

#### Brain parcellation

For each subject, the brain was parcellated in native DTI space into 90 different cortical and subcortical regions using the Automated Anatomical labelling (AAL) template^31^, where each region represents a node in the brain network.

Flirt (FMRIB, Oxford)^32^ was used to linearly co-register the standard ICBM152 in MNI space^33^ to each native T1-weighted structural image of each subject, using an affine registration (12 DOF) combined with a nearest-neighbour interpolation method. The resulting transformation was further applied to warp the Automated Anatomical Labeling (AAL). The MRI-DTI scan was then converted to MRI-T1 space, using a rigid-body transformation (6 DOF) and the resulting transformation matrix inverted. The MNI to MRI-T1 and MRI-T1 to MRI-DTI transformation matrices were subsequently concatenated, allowing a direct co-registration of the AAL template in MNI space to the diffusion MRI native space. This last transformation was performed using a nearest-neighbour interpolation method to ensure that discrete labelling values were preserved.

#### Brain structural network

We used the FDT toolbox in FSL (version 5.0, http://www.fmrib.ox.ac.uk/fsl/, FMRIB, Oxford) to carry out the multiple processing stages of the diffusion MRI data. The initial pre-processing involved correction of head motion and eddy current gradient induced image distortions. Subsequently, we modelled for crossing fibres within each voxel of the brain using a Markov Chain Monte Carlo sampling algorithm to build up distributions on diffusion parameters and estimate the local probability distribution of fibre direction at each voxel of the brain^34^. For this step, we used an automatic estimation of two fibre directions within each voxel, in order to improve the tracking sensitivity of non-dominant fibre populations in the human brain^35^.

Probability of connectivity was estimated using probabilistic tractography at the voxel level, with a sampling number of streamline fibres per voxel set at 5000. Brain boundaries were defined based on a binary brain previously registered in a skull-extracted version of the subject’s native brain. The connectivity from a seed voxel *i* to another voxel *j* was defined by the proportion of streamlines leaving voxel *i* and reaching voxel *j*^35^. This was then extended from the voxel to the region level, i.e. in a brain region consisting of *n* voxels, 5000\**n* fibres were sampled. The connectivity probability *P*_ij_ from region *i* to region *j* is then calculated as the number of sampled fibres in region *i* connecting the two regions, divided by 5000\**n*, where *n* is the number of voxels in region *i*. For each brain region, the connectivity probability to each of the remaining 89 AAL regions was calculated. The regional connectivity probability was computed using in-house Perl scripts and further normalised by each region’s volume, expressed in number of voxels. It should be noted that, given the dependence of tractography on the seeding location, the probability from *i* to *j* is not necessarily the same to that from *j* to *i*. However, these two probabilities are highly correlated across the brain for all participants. We therefore defined the undirectional connectivity probability *P*_ij_ between regions *i* and *j* by averaging these two probabilities. We considered this as a measure of the structural connectivity between each pair of areas, with *C*_ij_ = *C*_ji_.

For each participant, a 90×90 symmetric weighted matrix *C* was constructed, representing the brain’s structural network.

### Graph Theorethical Analysis

#### Standard Graph Metrics

The structural brain networks, represented as a 90×90 connectivity matrix, can be analysed as graphs. Using the Brain Connectivity Toolbox^36^, the brain networks were characterized using measures from graph theory. A rescaled version of the structural networks, where individual networks are normalised by its maximum connectivity value, was used. The following standard local and global graph metrics were considered: Connection Density, Node Degree, Local Efficiency, Global Efficiency, Characteristic Path length, Clustering Coefficient and Small-World index. Definitions and equations for these measures can be found in Supplementary Information. Additionally, the following topological features were assessed for both populations.

#### Modularity

Modularity quantifies the degree to which the network may be subdivided into non-overlapping communities. Therefore, the optimal community structure is represented as a subdivision of the network into well-defined groups of nodes so that it maximizes the number of within-group edges and minimizes the number of between-group edges. The calculation of the modularity coefficient was determined using the Louvain algorithm^37^. To circumvent the probabilistic nature of this algorithm, the optimal partition was selected from an iterative set of 1000 partitions, based on modularity maximisation statistic, i.e. the degree to which the network may be subdivided into clearly delineated groups.

#### Hubs

In this study, network hubs are defined as the most efficient nodes in a network, i.e. those with the highest *E*_*nodal*_. For each node *i*, if the normalised (divided by the mean *E*_*nodal*_ of all nodes) *E*_*nodal*_*(i)* is larger than the normalised mean *E*_*nodal*_ of all nodes in a network plus one standard deviation (SD), the node is considered a hub region^38^.

In addition to the hub classification based on *E*_*nodal*_, a set of hubs were classified by analysis of the distribution of connections within and between modules, allowing the categorization of hub regions into provincial and connector hubs, as described below.

*Provincial hubs* were defined as the nodes with a high within-module degree centrality (within module z-score greater than the mean plus SD of all nodes), indicating its central role in intra-modular communication, and low participation coefficient (PC) (PC≤0.3). Participation coefficient or proportion of cross-module connectivity profile compares the number of links of a given node to other nodes in different communities, to the total number of links to other nodes in the same community. This measure is defined as:

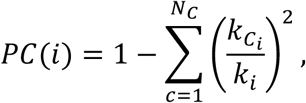

*Connector hubs* were defined as regions with a high value of within-module degree centrality and participation coefficient (PC>0.3), indicating a high proportion of cross-module connectivity, and thus a central role in the inter-modular communication. Upon identifying connector-hubs, we further characterized their connectivity to the rest of the network.

### Rich club

By definition, a set of brain areas in the network shows a rich club structure if its level of interconnectivity exceeds the level of connectivity expected on basis of chance alone^36^. In particular, the weighted rich club coefficient Φ*(k*) is computed as the ratio between the weights of connections present within the subnetwork *S*, composed of regions with a degree > k, and the total sum of weights present within an equally sized subset of the top ranking connection-weights in the network. The normalized rich club coefficient Φ_*norm*_(*k*) is computed by dividing Φ*(k*) by Φ_*random*_*(k*), with Φ_*random*_*(k*) calculated as the average rich club coefficient for each *k* of a set of 1000 randomized graphs (preserving the degree distribution)^36^. Therefore, a structural brain network can be described as having a rich club organization if, for a given degree interval, the normalized rich club coefficient is greater than 1.

The integrative nature of rich club members by the linking of different communities was also assessed. For this, we identified “rich-connector-hub networks”. These were constructed by examining the connectivity profile of nodes that simultaneously qualified as rich-club members and connector hubs (as defined above), to the remaining connector hubs in the network.

### Between-group differences in structural connectivity strength

Between-group statistical comparison of the raw structural connectomes for PBD patients and controls was performed on a connection-by-connection basis, using a non-parametric statistical method – network-based statistics (NBS). This method allows the identification of significantly altered sub-networks, while controlling for the family-wise error rate (FWER)^39^. A data exploration step determined the t-threshold for this specific dataset, consisting of analysing the distribution of node count of the largest component versus a large range of t-thresholds. Based on statistical conservativeness (maximal t-threshold where a component was found), a primary statistical threshold of *t*=2.1 (Supplementary Figure 1). Recent findings of abnormal brain networks schizophrenia suggest the same value as an optimal t-threshold, based on the criteria of maximal node count provided by a t-threshold ≥ 2.00^40^. The topological configuration of a group effect in structural connectivity strength was then represented by one or multiple significantly connected components with FWER-corrected *p*<*0.05*.

### Between-group statistical differences in Graph measures

Between-group statistical comparison was performed using M-W U test, KS-test and T-test.

### Fingerprints of modular connectivity – within- and between-group analysis

In order to investigate potential alterations in modular connectivity of PBD with psychosis, we analysed the network fingerprints of inter-modular (global and connector-hub-driven) and intra-modular structural connectivity, for each group. Modular connectivity strength was quantified as the total number of connections (degree) of all nodes forming a module. In order to allow for group comparison, a single community structure was used as the reference scheme to quantify modular connectivity for both populations. The criteria for selecting the reference scheme was based on which population revealed a higher mean score of community-structure goodness-of-fit, i.e. how well the estimated group community structure represents each subject’s individual community structure. Maps of ‘between-module connectivity’, ‘within-module connectivity’ and ‘connector-hub modular connectivity’ were then produced for each of the two populations.

The topological characterisation of each module was done by computing the mean degree, participation coefficient, betweeness-centrality and within-module degree centrality of all nodes in a module’s network. Between-group differences in modular network topology were investigated using non-parametric statistical testing (permutation-based unpaired t-tests). Correction for multiple comparisons was performed using FDR-correction.

### Relationship between network metrics and neurocognitive scores

The neurocognitive scores (coding and FSIQ) assessed for the HC and PBD groups, as well as the positive and negative symptoms scores (PANSS) assessed in the PBD patients, were correlated with local and global network properties. Partial correlation analysis was performed, using age and gender as confounding variables. Similarly, the correlation between clinical scores and the subset of network nodes revealing significant group differences in nodal efficiency were also assessed using age and gender as confounding variables.

## Results

### Demographics

No significant differences were found for age (*p*=0.30), gender (*p*=1), verbal IQ (*p*=0.20), performance IQ (*p*=0.16) and FSIQ (*p*=0.13) scores between patients with Pediatric Bipolar Disorder (PBD) and healthy controls (HC) (Table 1).

### Connectivity Strength

Compared to controls, patients with PBD showed significantly altered structural connectivity (*p*=0.015) in a brain sub-network, or connected component, involving 71 structural links (~1,8% of all possible connections; ~5,5% of all existing connections) (Figure 1). These 71 links showed 39 decreases and 32 increases in connectivity strength in the PBD group (see Supplementary Table 1 for details). The network is predominantly distributed in right frontal inferior and temporal brain areas (22% more connections on the right hemisphere).

**Figure 1.**
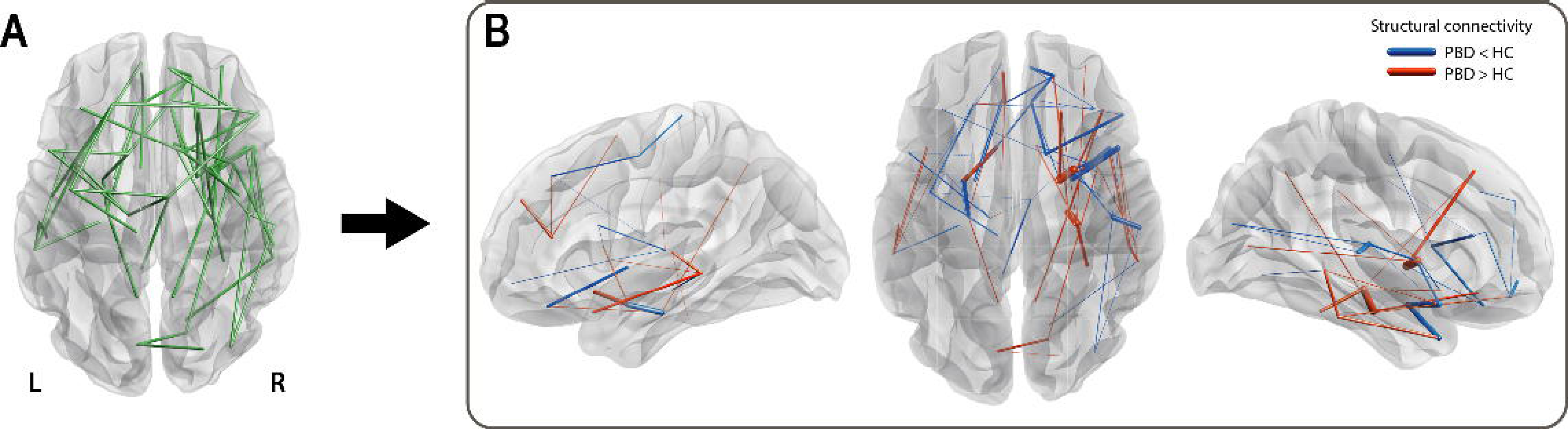
Significant changes in the structural connectivity of patients with pediatric bipolar disorder (PBD) with psychosis, compared to healthy controls (HC). Connected component of significant differences in structural connectivity strength between patients with PBD and HC, represented with edges connecting a pair of regions. A) Binarised version of the connected network of significantly altered structural connectivity in PBD. B) Weighted version of A), where the edge thickness represents the amplitude of the differences in structural connectivity. Decreases and increases in connectivity strength between regions in the PBD group are represented in blue and red respectively. PBD primarily differs from HC in inferior temporal networks (e.g. left and right temporal pole to left and right amygdala, left amygdala to left parahippocampal region) and inferior frontal networks (e.g. right orbitofrontal cortex) and subcortical to inferior frontal regions (e.g. right caudate to orbitofrontal cortex and left pallidum to left orbitofrontal cortex).

### Graph Theoretical Analysis: Local Properties

#### Nodal efficiency - network hubs

For each group, regions showing a high normalized nodal efficiency were identified as network hubs. Between-group differences found in the network hub configuration are suggestive of local asymmetries in efficiency of communication (Figure 2).

**Figure 2.**
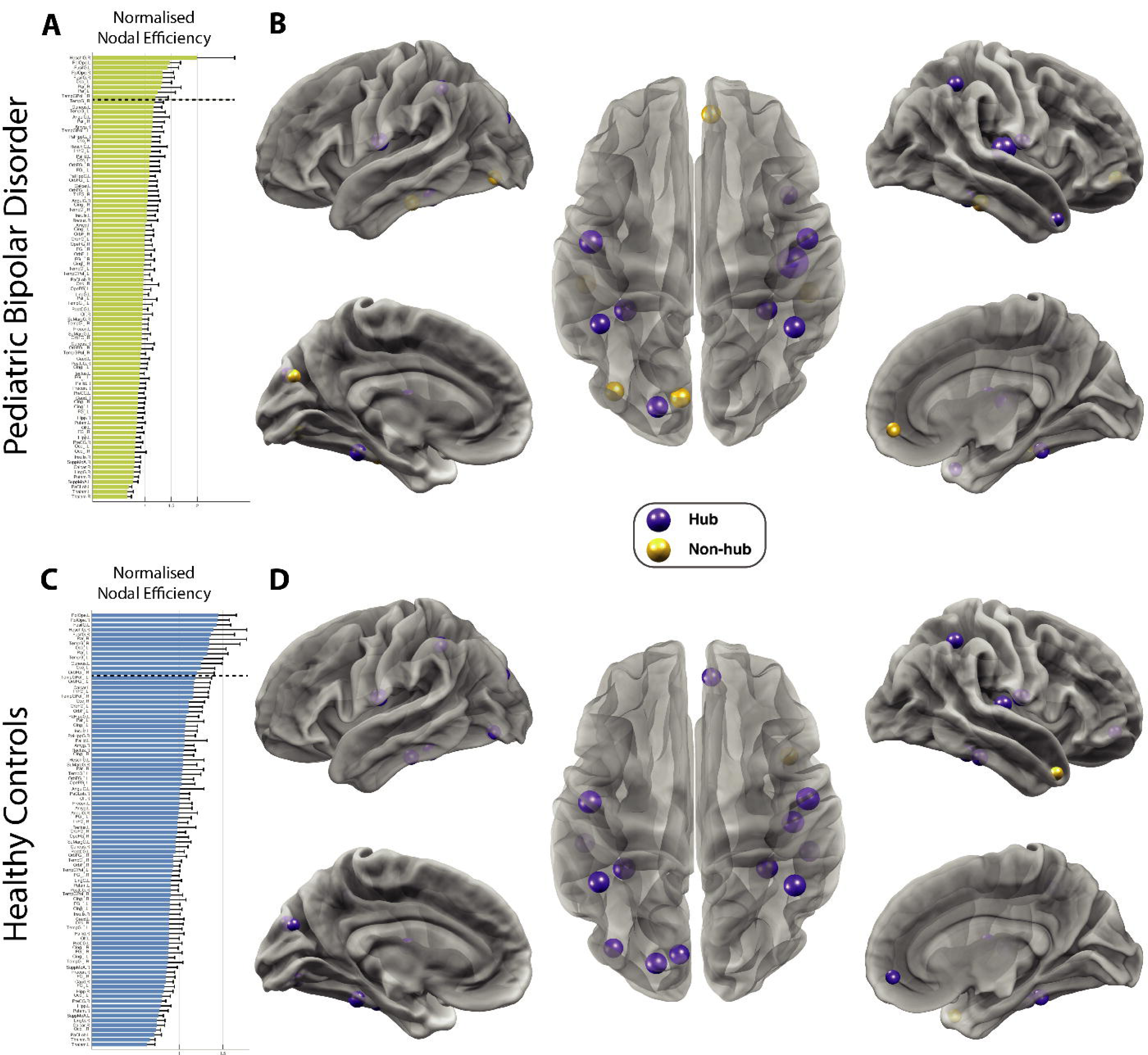
Changes in network hub regions in patients with pediatric bipolar disorder (PBD) wth psychosis. Differences in network hub regions in HC (top row) and PBD (bottom row) as measured by the normalised nodal efficiency for all 90 AAL brain regions. This shows a clear reorganisation in hub regions in PBD where e.g. the orbitofrontal cortex is no longer a hub region,. **A)** For the patient group, the figure shows the nodal efficiency sorted according to the mean in descending order. For each node, the *E*_*nodal*_ is normalised by the mean of all nodes’ *E*_*nodal*_, and a node is identified as a hub region if its normalised *E*_*nodal*_ is larger than the sum of the mean plus the SD of all network nodes’ *E*_*nodal*_. **B)** Hub regions, represented as spheres positioned according to the centroid stereotaxic coordinates of the correspondent anatomical region, with node size proportional to their *E*_*nodal*_, are mapped onto a 3D reconstructed brain surface. **C)** The ranking of nodal efficiency, and **D)** the identified hub regions for healthy participants. For each group, the existing hub nodes are represented in purple and non-hub nodes (hubs exclusive to the other group) in yellow.

In the HC group, 13 hub areas were identified (Figure 2a; Supplementary Table 2). The hub configuration for the PBD group revealed the loss of five hubs and inclusion of one new hub when compared to the HC group. Of particular interest are the exclusivity of the medial orbitofrontal and inferior temporal (bilateral) hubs to healthy controls.

### Graph Theoretical Analysis: Global properties

No significant network differences were found between the HC and PBD structural connectomes on any of the global graph measures assessed: global efficiency, modularity, clustering coefficient, characteristic path length and small-worldness (see Supplementary Table 5).

### Modularity and Hub Classification

For each group, the whole-brain optimal community structure was decomposed into seven modules as shown in Figure 3A. The HC and PBD groups revealed differences in their modular arrangement, with the most important changes found in the right lateral and the medial posterior regions of the brain.

**Figure 3.**
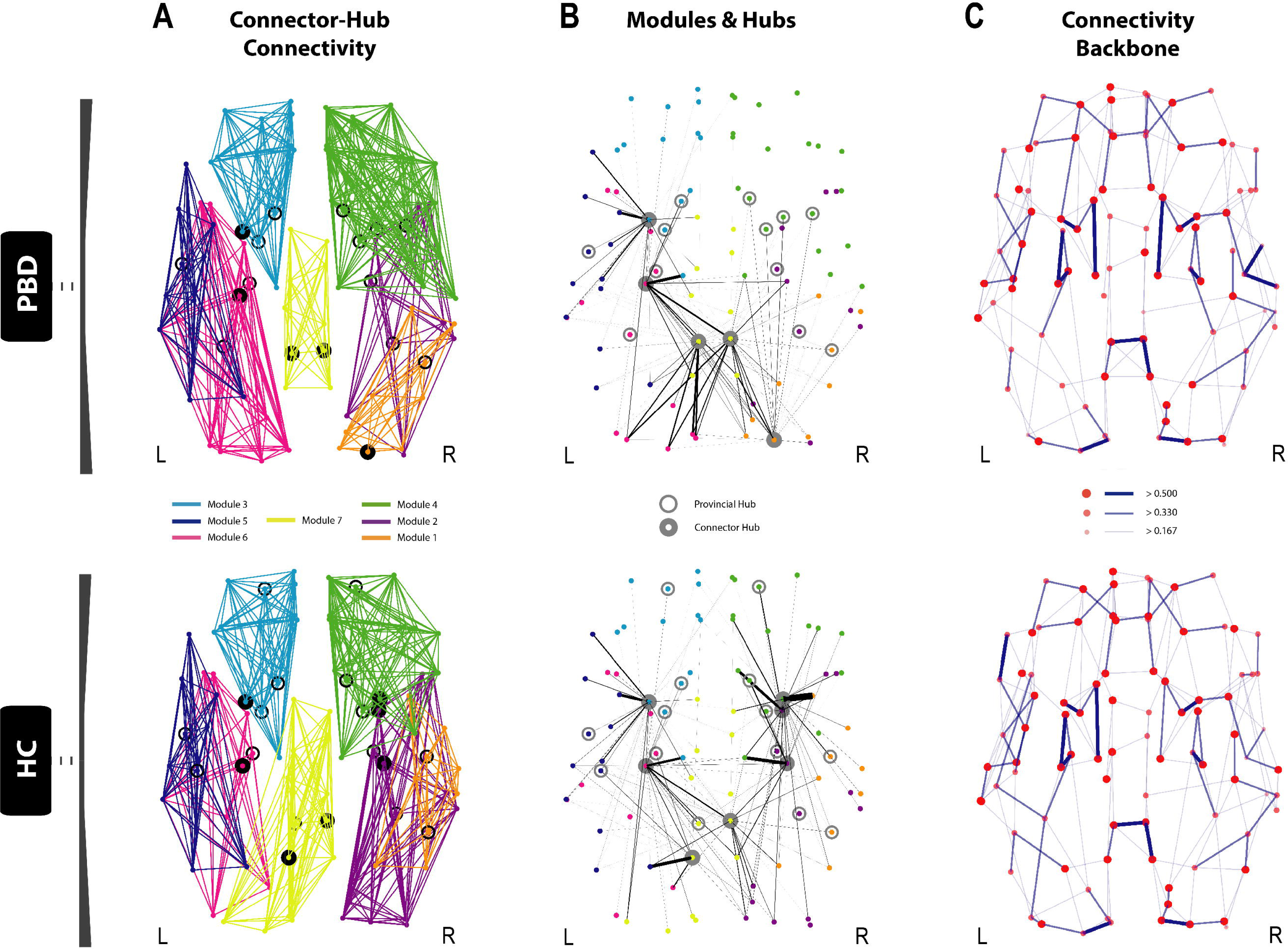
Disrupted modularity, hubs and connector-hub connectivity in patients with pediatric bipolar disorder with psychosis. The figure shows the changes in the optimal community structure, i.e. the seven modules and connector hubs shown on a dorsal view of the brain of patients with PBD (top row) and healthy controls (HC, bottom row), with node colours denoting membership to a module and with intra-module edges coloured accordingly. *Connector hubs* (marked as filled circles) are defined as regions with high within-module degree centrality and a participation coefficient p>0.3, denoting a high proportion of cross-module connectivity. *Provincial hubs* (marked as unfilled circles) are defined as having high within-module degree centrality but participation coefficient p≤0.3. **A)** Dorsal view of the optimal modularity partitioning found in PBD (top), reveals a different configuration compared to HC (bottom). **B)** Significant local changes are found in the undirected structural connectivity profile for the connector hubs in PBD (top) compared to HC (bottom). Edges, coloured in black, correspond to the structural connectivity of each connector hub region to any other region in the brain, with thickness proportional to its connection strength. Colour of dots represents the belonging of a brain area to a particular community, consistent with A). Albeit the number of both provincial and connector hubs is preserved between the two groups (Supplementary Table 2), variance in the distribution reflects the high impact on the regional dispersion and density of connector-hub-connectivity (CHC), i.e. network of undirected connections involving connector hubs. Whilst there is an almost symmetric distribution of provincial and connector hubs in the HC group, asymmetries found in the hub distribution in the PBD group led to a significantly decreased CHC density in the right anterior hemisphere and a shift of the CHC towards posterior areas of the brain, as shown in **B)**. Notably, the connection between the orbitofrontal provincial hub and the amygdala connector hub is missing in the patient group. C) Dorsal view of the brain connectivity backbone for patients with PBD with psychosis (top), and HC (bottom). Nodes and edges are coded according to normalised degree and connection weight respectively. Diameter, thickness and transparency of nodes and edges are divided into three weight levels (see legend).

#### Rich club

Rich club organisation was found in the structural networks of both HC and PBD patients. The group-weighted and normalised rich club coefficient curves of both HC and PBD, shows that the maximal Φ_*norm*_(*k*) is reached at *k*=35 for both groups. However, as illustrated in Figure 5A, the peak amplitude is clearly decreased in the patient group, which is indicative of a reduced rich club organisation for this group. This reveals lower level of connectivity between the most densely connected regions in the brain, when compared with the HC group. This effect appears clear and reflects the interval of largest difference between the two curves. However, given that it only extends for a limited range of *k* values (34≤*k*<37), it is not sufficiently large to reject the null hypothesis of no difference between means. Additionally, rich club membership between groups differed only in two right hemisphere nodes, suggesting that the insular cortices and the hippocampus have a rich club profile exclusive to the PBD group (Figure 5B).

As described in Methods section *Rich Club*, we also examined the connectivity profile of nodes that simultaneously qualified as rich-club members and connector hubs to the other connector hubs in the network (Figure 5C). These networks revealed a relatively symmetric connectivity pattern and central spatial distribution in the HC group, involving regions of the right posterior cingulum and amygdala, left precuneus and left and right hippocampus and putamen (Figure 5C (bottom)). This strong ‘core-effect’, supported by a structural network of rich-connector-hubs linking functionally different communities, reflects a tendency toward efficient integration of information across the brain, in the HC group. In contrast, the PBD group exhibited a comparatively asymmetric profile of ‘rich-connector-hub’ connectivity (Figure 5C (top)). This tendency toward spatially unbalanced modular integration was characterised by a deficit of two right hemisphere rich-connector-hubs (loss of the hippocampus, amygdala and putamen; gain of the superior occipital gyrus), when compared to the HC group, resulting in a single right hemisphere module (posterior) being comprised in this network, as opposed to two right-hemisphere modules in the HC group.

### Modular Connectivity

Our findings suggest that there are relevant alterations in the fingerprints of structural inter-modular connectivity in PBD (Figure 4). Specifically, in PBD, the connectivity of module 4 (right hemisphere; thalamus, insula, basal ganglia, anterior cingulate, precentral gyrus and regions covering most of the frontal cortex) is increased to module 1 (right hemisphere; amygdala, hippocampus, parahippocampus, fusiform gyrus, and other regions of the occipital and temporal cortices) and decreased to module 3 (left hemisphere; thalamus, basal ganglia, anterior cingulate and a large extension of the frontal cortex). Overall, module 1 has overall increased inter-modular connectivity but decreased connector-hub driven inter-modular connectivity, which suggests that PBD is associated to an increased degree of sparsity in inter-modular linkage (i.e. more connections, but a less prominent role of its connector-hubs in mediating inter-modular connectivity). Additionally, despite the absence of connector-hubs in module 4 in PBD, its mean inter-modular connectivity strength is the second largest across all modules and remains approximately the same as in HC (with two connector-hubs). This may suggest that more nodes (with no ‘hub’ or ‘connector’ profile – lower degree and/or lower participation coefficient) are mediating communication between module 4 and all other modules in PBD, which may be indicative of a decreased efficiency/selectivity in communication processes between a network including left hemisphere limbic, insular and thalamic areas, and the rest of the brain.

**Figure 4.**
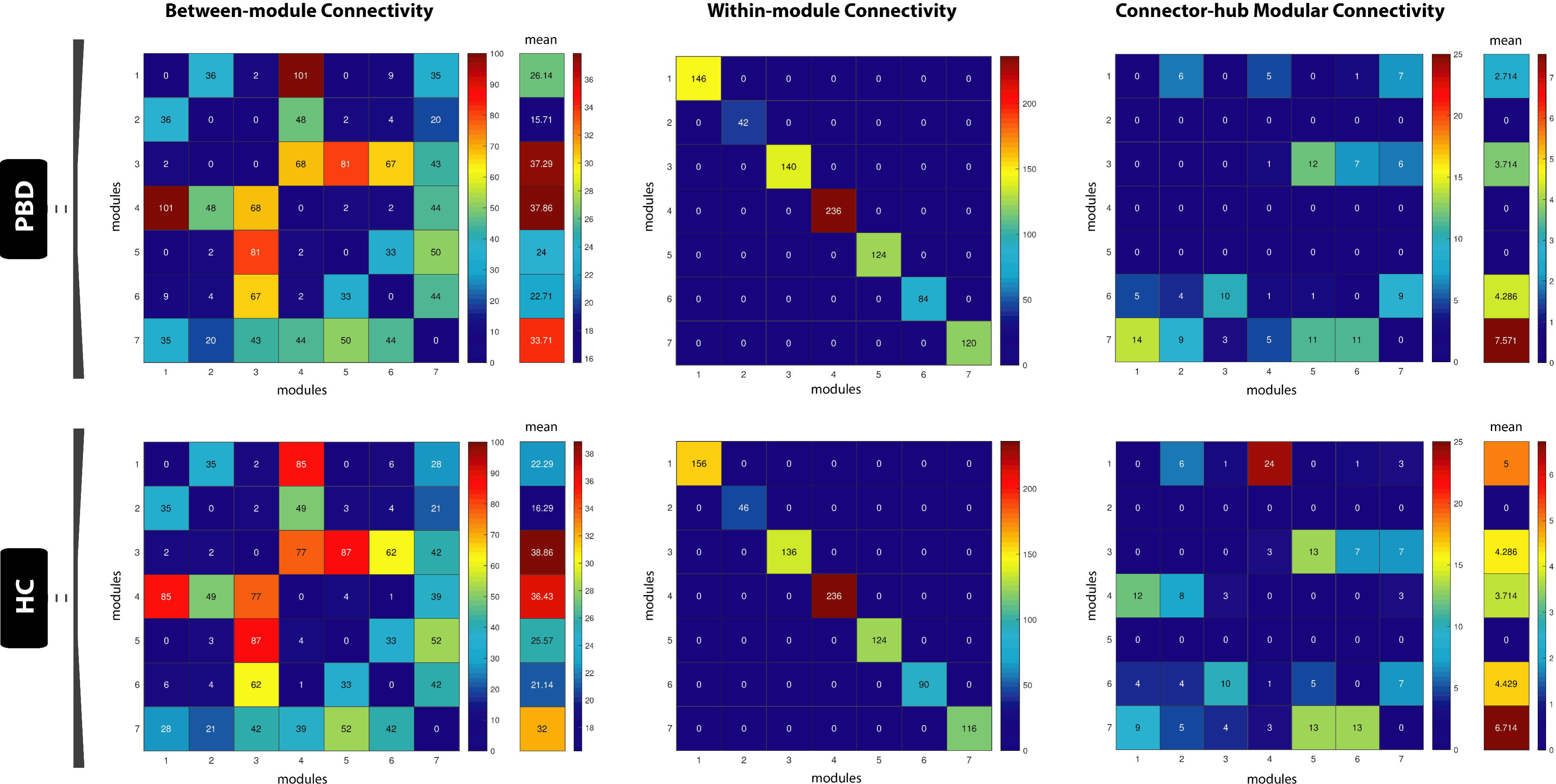
Modular network analysis. The figure shows the patterns of inter-modular (left column), intra-modular (middle column) and connector-hub driven inter-modular (right column) connectivity, for patients with PBD with psychosis (top row) and HC (bottom row). Modular connectivity strength is quantified as the total number of connections (degree) of all nodes forming a module. The resulting community structure for the HC population was selected as the reference scheme for group modular connectivity analysis, given its higher group goodness-of-fit (i.e. how well the group estimated community structure fits to each subject’s individual community structure). The figure shows that the patterns of SC between modules are changed in PBD. Specifically, that the connectivity of module 4 (right hemisphere; thalamus, insula, basal ganglia, anterior cingulate, precentral gyrus and regions covering most of the frontal cortex) is increased to module 1 (right hemisphere; amygdala, hippocampus, parahippocampus, fusiform gyrus, and other regions of the occipital and temporal cortices) and decreased to module 3 (left hemisphere; thalamus, basal ganglia, anterior cingulate and a large extension of the frontal cortex), whereas module 1 has overall increased inter-modular connectivity but decreased connector-hub driven inter-modular connectivity. The latter suggests an increased degree of sparsity in inter-modular linkage for module 1 (i.e. more connections, though a less prominent role of its connector-hubs in mediating inter-modular connectivity). Despite the absence of connector-hubs in module 4 in PBD, its mean inter-modular connectivity strength is the second largest across modules and similar to HC (with two connector-hubs). This suggests that more nodes (with no ‘hub’ or ‘connector’ profile) are mediating communication between module 4 and the remaining modules in PBD. This may be indicative of a decreased efficiency/selectivity in processing communication between a network including the left hemisphere limbic, insular and thalamic areas, and the rest of the brain.

**Figure 5.**
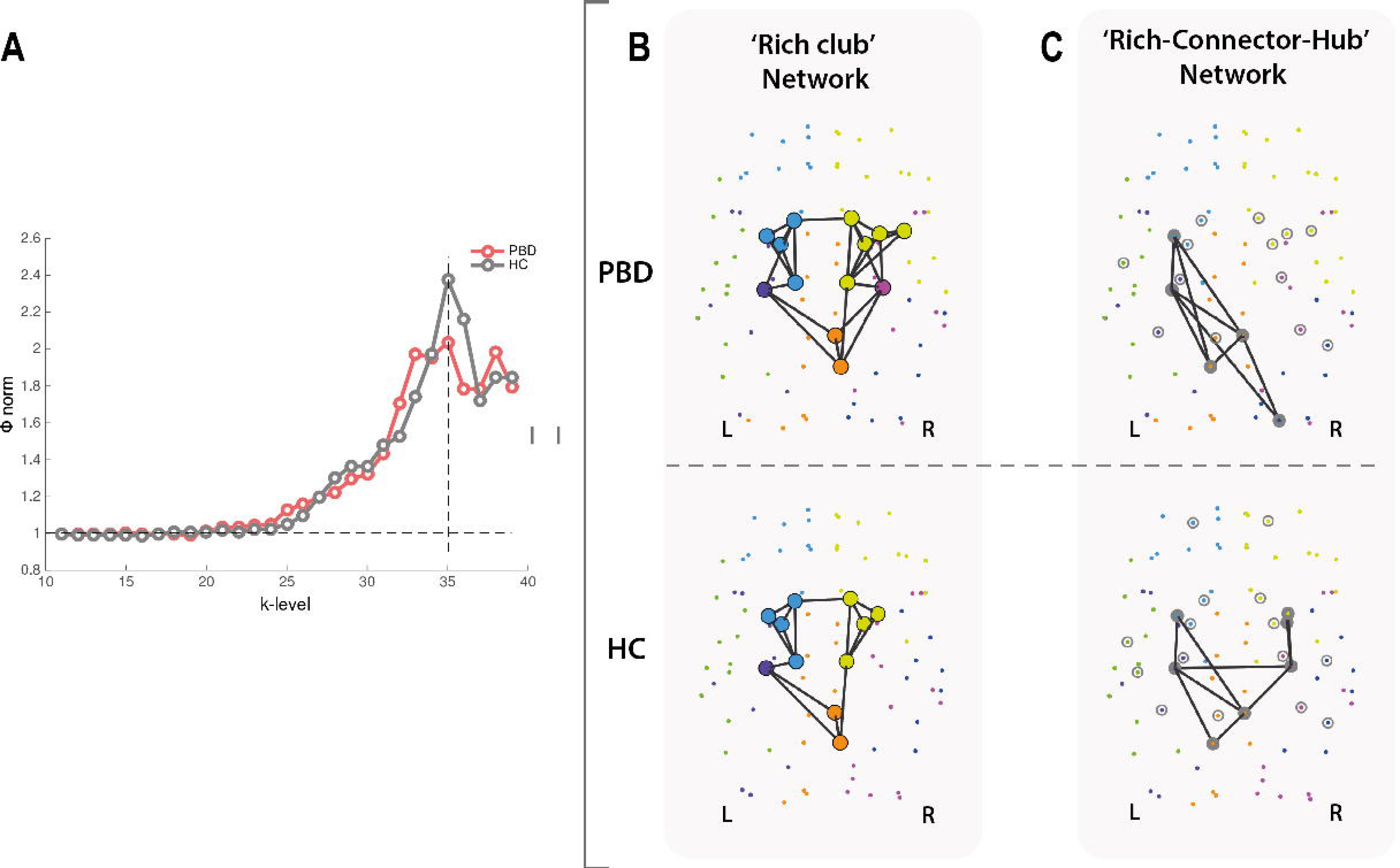
Changes in rich club organisation and inter-modular integration in pediatric bipolar disorder (PBD). **A)** The figure shows significant changes in the rich club organisation as demonstrated by the normalised weighted rich club coefficient as a function of *k*-level (degree), for the PBD (red curve) and healthy control (HC, grey curve) groups. **B)** Rich-club members in HC (bottom) and PBD (top). Rich club nodes are represented by large filled circles with colours denoting the module to which they belong. Rich club member regions common to both groups included the left hippocampus, right posterior cingulate cortex and precuneus, and left and right caudate, putamen, pallidum and thalamus. Membership between groups differed only in two right hemisphere nodes, suggesting that the insular cortices and the hippocampus have a rich club profile exclusive to the PBD group. **C)** Equally, the ‘rich-connector-hub’ network signature, i.e. the backbone of inter-modular connectivity driven by connector-hubs, is differed between the HC (top) and PBD (bottom). Edges coloured in black represent the existing structural connectivity between connector hubs.

Significant group differences in modular network topology were found in degree (module 2 and 4), within-module degree centrality (module 7) and participation coefficient (module 6) (table 2). However, only the last significant finding (module 6 – participation coefficient) survived to correction for multiple comparisons.

### Network properties vs. neurocognitive and psychotic symptoms

We examined the potential association of neurocognitive scores – coding, VIQ, PIQ, FSIQ – and psychotic symptoms with significant group differences in nodal efficiency (Supplementary Table 3), as well as with other global graph properties (Supplementary Table 4).

It is only for the orbitofrontal cortex that nodal efficiency was significantly correlated with IQ and/or psychotic symptoms (PANSS). Nodal efficiency of the left inferior orbitofrontal cortex was found positively correlated with the VIQ (r=0.74; *p*=0.04), PIQ (r=0.60; *p*=0.03) and FSIQ (r=0.77; *p*=0.02) scores, whereas negative psychotic symptoms were positively correlated with the nodal efficiency of the left middle orbitofrontal cortex (r=0.57; *p*=0.04) (Figure 6). These findings however did not survive correction for multiple comparisons.

**Figure 6.**
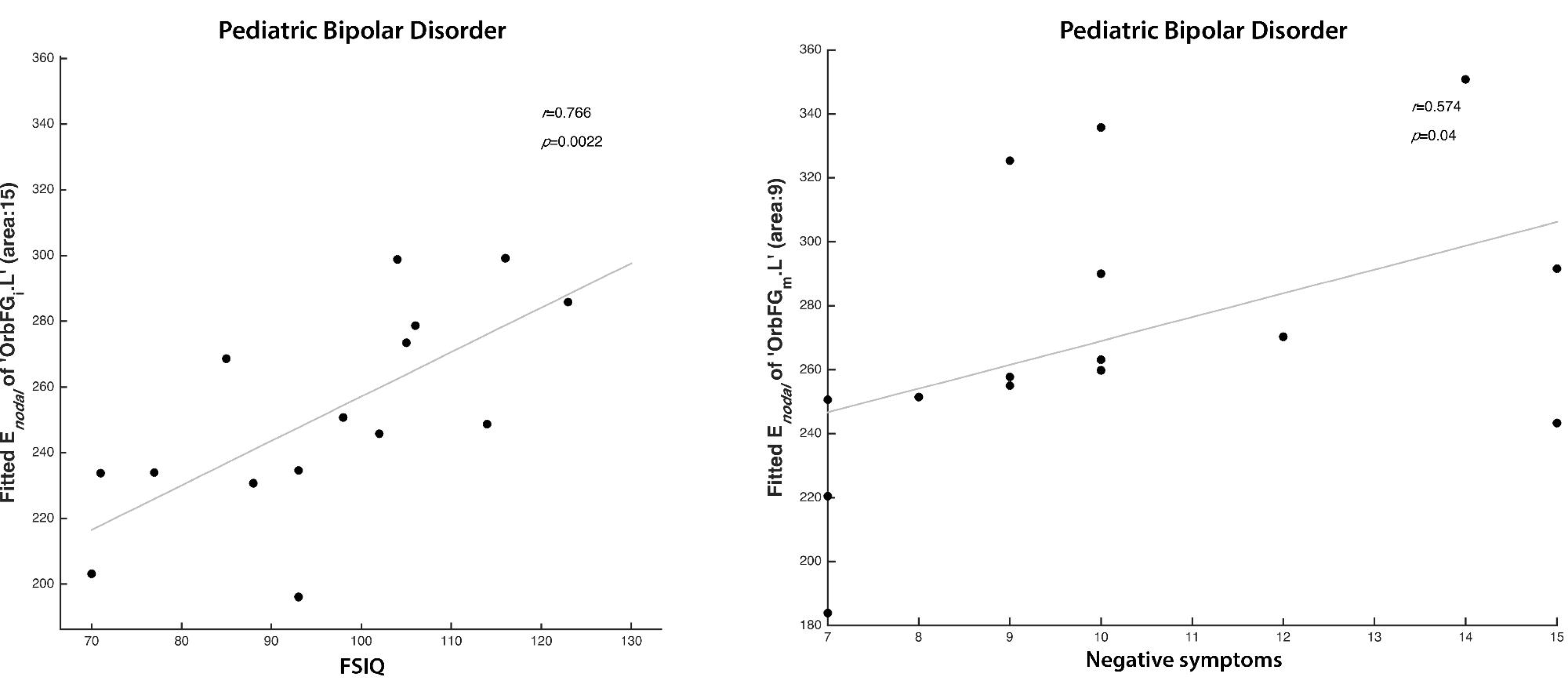
Nodal efficiency of the orbitofrontal cortex in a whole-brain structural connectivity network correlates with psychotic and IQ scores of patients with pediatric bipolar disorder (PBD). **Left:** A significant positive correlation is found between the nodal efficiency of left inferior orbitofrontal cortex and FSIQ. **Right:** A significant positive correlation is also found between the nodal efficiency of the middle orbitofrontal cortex and negative psychotic symptoms in the PBD group assessed with the PANSS (see Supplementary Table 3 for full details).

None of the global network metrics assessed showed significant correlation with the neurocognitive/psychotic scores (Supplementary Table 4).

## Discussion

We investigated the topological organisation of the structural connectomes in a cohort of adolescents with a diagnosis of PBD with psychosis. Network analysis using a combination of state-of-the-art methods revealed significant differences in structural connectivity between PBD patients and healthy matched participants. Most differences were found in networks involved in emotional regulation, strengthening the hypothesis that BD could be linked to prefrontal top-down dysregulation of emotional processing. Specifically, we found widespread differences in structural connectivity strength, and altered hub configuration and connectivity profile. Changes were also found in the rich-club organisation of the PBD structural network as well as in inter-modular connectivity driven by ‘rich-connector-hubs’.

These findings reveal that key network features for balanced whole-brain integration of information are altered in PBD. In addition, our results suggest that the nodal efficiency of regions important for the regulation of emotional processing such as the orbitofrontal cortex show significant positive correlations with IQ score and negative symptoms in PBD patients. Our results are in line with recent evidence of association between altered structural connectivity in a cortico-striatal reward circuit, particularly involving the OFC, and hypomania symptoms^41^. Overall, these findings of abnormal topological organization in PBD shed new light on the potential neurobiological deficits underlying BD.

The widespread alterations in structural connectivity in the PBD brain networks (shown in Figure 1), primarily differ from HC in inferior temporal networks and inferior frontal networks and subcortical to inferior frontal regions (Figure 1; Supplementary Table 1). This fits well with previously reported localised changes in white matter, where there is evidence for a loss of connectivity involving prefrontal and frontal regions through associative and commissural fibres^42,43^. Widespread *decreases* in fractional anisotropy (FA) have been found in major tracts, including (but not limited to) the orbitofrontal cortex^20^, superior frontal lobes^16,43^, and bilateral parietal and occipital corona radiate^19^. It has indeed been suggested that white matter changes could be central to BD, and may represent an endophenotype^44^. Our findings of widespread connectivity changes in PBD support this idea and add significant information on the specific structural connectivity networks and brain topological features that are affected.

Our structural connectivity findings are also consistent with the previously reported widespread changes in regional grey matter volumes in PBD. Specifically, neuroimaging studies have linked PBD^45^ to altered cortical grey matter volumes in several regions shown to have abnormal structural connectivity in the present study, including (but not limited to) the basal ganglia^46^, thalamus^46^ and temporal cortices^46,47^, cingulate^45–48^, dorsolateral prefrontal cortex (DLPFC)^49,50^, temporal lobe^2,46,51^, orbitofrontal cortex (OFC)^46,49,2,50^, and the amygdala. The substantial overlap between previously reported grey matter changes and the structural connectivity changes demonstrated here, are suggestive of a mechanistic link between the volume of cortical and subcortical regions involved in emotional regulation and their structural connectivity to the rest of the brain.

Furthermore, our structural connectivity findings can be thought of in terms of providing the supporting connectome for the previously observed *functional* connectivity changes in PBD. The structural changes in PBD are likely to lead to changes in functional brain activity. Thus far few studies investigating resting state functional connectivity in BD have been published, but changes in the functional connectivity of the orbitofrontal cortex^2,46,52^, amygdala^53^ and temporal lobe^46,54^ have been reported. One particular study analysed resting state networks in BD during the manic state and showed significant changes in resting state connectivity, mainly involving regions associated with the fronto-temporal, ventral-affective and dorsal-cognitive circuits^55^. Recent evidence has linked BD to a decreased structural connectivity in right hemisphere in a sub-network involving connections between fronto-temporal and temporal areas^23,29^. Additionally, self-report measure of bipolar risk (hypo/mania proneness) has been linked to increased connectivity between the nucleus accumbens (part of the Caudate in the AAL parcellation), amygdala and mOFC^41^. These observations are consistent with the present structural connectivity findings (e.g. temporal pole to amygdala, amygdala to caudate and parahippocampal gyrus, superior frontal gyrus and pallidum, between orbitofrontal cortices, caudate and pallidum to orbitofrontal cortices). Taken together, our findings strengthen the hypothesis that functional network dysregulation supported by abnormal structural connectivity, is a prominent driver of pathophysiology in PBD.

The results from our graph theoretical analysis of the PBD connectome also revealed significant changes on the nodal efficiency measure with a loss of five hubs (right orbitofrontal cortex, left cuneus, bilateral temporal gyrus and left occipital cortex) in PBD relative to HC. These brain regions have been demonstrated to be involved in emotional processing and regulation. As such, these changes are consistent with the previously reported abnormal prefrontal top-down dysregulation in BP^4,56^.

We further investigated the whole-brain optimal community structure for the PBD and HC. The networks were decomposed into seven modules and we found that the healthy controls and PBD groups showed differences in modular arrangement, as well as in the fingerprints of inter-modular connectivity. In PDB, a large community of brain regions covering most of the right frontal lobe (including the thalamus, insula, basal ganglia, anterior cingulate cortex, amongst other regions), showed increased intra-hemispheric connectivity to an occipito-temporal module (comprising regions such as the amygdala, para/hippocampus, and fusiform gyrus), and decreased inter-hemispheric connectivity to a large module in the left frontal lobe. Additionally, our results suggest that PBD is associated to a decreased degree of centrality in inter-modular connectivity, given a weaker role or inexistence of connector-hubs in mediating inter-modular connectivity, i.e. despite the total number of connections linking modules is similar in PBD, these are mostly driven by a higher number of nodes with a less central role – non-hubs. This effect is found in two right-hemisphere modules – frontal and occipito-temporal, and may be indicative of decreased efficiency in integrating information between these modules and the rest of the brain. Furthermore, significant group differences in participation coefficient were found in a left occipito-temporal module.

The potential functional relevance of these modularity changes can notably be further characterised by investigating the rich club organization. Interestingly, at peak amplitude k=35 for the HC, we found the degree of rich club organisation in PBD to be lower than in HC. This considerable reduction in rich club organisation suggests lower connectivity between topologically central hub regions of the brain, compared to the healthy participants. To further characterize these changes, we analysed the networks of structural inter-modular connectivity that are driven by ‘rich-connector-hubs’; that is rich-club members that also qualified as connector hubs with high betweeness-centrality and high participation coefficient. Despite the considerable level of overlap in the rich club membership between groups, the integrative nature (participation in inter-modular connectivity) of rich-club members revealed a clear difference in the spatial distribution and number of involved modules, which is suggestive of a disruptive neural network integration capability in the PBD group. Specifically, a single right hemisphere module (posterior) was comprised in the rich-connector-hub network, which may reflect a tendency toward weaker global network integration in the PBD group.

We also demonstrated that the nodal efficiency of two subdivisions of the orbitofrontal cortex, is significantly correlated with IQ scores and negative symptoms in PBD patients, consistent with prior studies involving the orbitofrontal cortex in emotional processing^57^. Furthermore, structural connectivity changes may result not only in the emotional dysfunction seen in PBD but also in significant cognitive deficits as revealed in a recent meta-analysis of verbal learning and memory, processing speed, and executive dysfunction^58^.

In summary, our results show that there are significant changes in structural connectivity between patients with early–onset bipolar disorder with psychosis compared to healthy matched participants. These changes are primarily manifested in reduced connectivity between inferior, frontal and temporal cortical areas and to regions linked to emotion, memory and executive function, such as amygdala, hippocampus and the basal ganglia. Taken together, these findings suggest that PBD with psychosis is characterised by a dysfunctional prefrontal regulatory mechanism in line with the hypothesis of prefrontal top-down dysregulation in BD^4,56^. Furthermore, differences in the structural network’s topological organisation suggest a significant reduction of overall integrative processing capability, which is associated with the patients’ neurocognitive faculties and symptoms. These changes may potentially explain the deficits in emotion processing and regulation found in PBD, and ultimately lead to the development of more targeted treatments.

## Acknowledgements

The authors gratefully thank Mikkel Petersen for his help in the DTI analysis. MLK was supported by the ERC Consolidator Grant: CAREGIVING (n. 615539), TrygFonden Charitable Foundation and by Center for Music in the Brain, funded by the Danish National Research Foundation (DNRF117). GD was supported by the ERC Advanced Grant: DYSTRUCTURE (n. 295129), by the Spanish Research Project SAF2010-16085 and the FP7-ICT BrainScales. The clinical and neuroimaging data used in this study was supported by the Medical Research Council (MRC) and the Oxford Hospital Services Research Committee (OHSCR).

## Data availability

The datasets generated during and/or analysed during the current study are available from the corresponding author on reasonable request.

## Contributions

HMF designed and implemented the methodological framework used in this study. HMF, CG and TVH processed the data. HMF, MLK, JC, PP and LDL wrote the manuscript. AJ performed the clinical assessment of all patients and healthy participants and provided the financial support for the neuroimaging data collection. AJ, GD, PCW and AM acted in an advisory and editorial capacity.

