## Supplementary Information for "Disrupted brain structural connectivity in Pediatric Bipolar Disorder with psychosis"

### *Graph Theoretical Metrics*

The following graph theoretical metdics were computed in the present study.

*Connection Density*. Connection density, or wiring cost, is the actual number of edges in the graph as a proportion of the total number of possible edges. For an undirected graph with $N$ nodes without self-connections, the total number of possible connections *is* $(N * (N - 1))/2$.

*Degree*. The degree of a node *d(n)* is calculated as the number of nodes to which node *n* is connected.

*Characteristic Path Length.* The shortest path length between areas *n* and *p* is calculated as the average shortest path length in the network, i.e. the minimum number of connections, or minimum cost needed to connect regions *n* and *p*, where a connection cost, or edge length, is the inverse of connection weight. The characteristic path length is given as the mean shortest path length over all pairs of nodes in the network, calculated as the global mean of the distance matrix.

*Global Efficiency*. Global efficiency, $E_{global}$, reflects how efficiently information can be exchanged over the network and is defined by the mean of the inverse shortest path length, $L_{ij}$, between each pair of nodes in a network, $C$, with $N$ nodes (52):

$$E_{global}(C) = \frac{1}{N(N-1)} \sum_{i \neq j \in C} \frac{1}{L_{ij}}$$

*Nodal Efficiency*. The nodal efficiency, $E_{nodal}(i)$, reflects how well a node connects to all other nodes in the network and is defined as the mean of the inverse shortest path length, $L_{ij}$, between a node, *i,* and all other nodes in the network:

$$E_{nodal}(i) = \frac{1}{N-1} \sum_{i \neq j \in C} \frac{1}{d_{ij}}$$

*Local Efficiency*. The local efficiency of a network, $E_{local}$, is defined as the average *Nodal Efficiency*, and indicates globally how information is transferred within the neighbours of a given node.

$$E_{local}(C) = \frac{1}{N} \sum_{i \in C} E_{nodal}(i)$$

*Clustering coefficient.* The clustering coefficient reflects the extent of local interconnectivity in a network by considering the fraction of a node's neighbours that are also neighbours of each other. It is measured as the fraction of connected triangles, $\delta_v$, to the total number of triples in the network, $\tau_v$. A network's weighted clustering coefficient is then defined as the average of the clustering coefficient over all nodes.

$$Cl(C) = \frac{1}{|V'|} \sum_{v \in V'} \frac{\delta_v}{\tau_v}$$

where $V'$ is the subset of nodes with degree >2.

*Small-worldness*. A network C is considered to be small-world ($\sigma$>1), if the average shortest path length, $L$, is small and the weighted clustering coefficient, $Cl$, is high, when compared to equivalent random networks ($L_R$ and $Cl_R$, respectively) (53).

$$\sigma\ (C) = \frac{Cl/Cl_R}{L/L_R}$$

In this study we generated 100 matched random networks, preserving the degree distribution and ensuring connectedness. The ratio between the original and the generated random networks, for both the weighted clustering coefficient and weighted path lengths measures, allows the correction for network differences across individuals with regards to edge number and degree distribution.
