## Supplementary Table for "Disrupted brain structural connectivity in Pediatric Bipolar Disorder with psychosis"

### Supplementary Tables

**Supp. Table 1. Connections comprising the connected component of significant structural connectivity difference between PBD and HC ($p < 0.05$).**

| Area1 | | | Area2 | | Difference |
|---|---|---|---|---|---|
| name | n | | name | n | |
| TempGPol_s (R) | 42 | <-> | Amyg (R) | 84 | -214.01 |
| Amyg (L) | 39 | <-> | PaHippG (L) | 41 | -204.23 |
| TempGPol_m (R) | 42 | <-> | Amyg (R) | 88 | -194.00 |
| TempG_s (R) | 80 | <-> | HeschlG (R) | 82 | -192.49 |
| OrbFG_m (R) | 6 | <-> | OrbF_s (R) | 26 | -123.43 |
| Pallid (L) | 27 | <-> | Rectus (L) | 75 | -119.22 |
| Caud (R) | 14 | <-> | TriFG_i (R) | 72 | -119.15 |
| Caud (R) | 28 | <-> | Rectus (R) | 72 | -103.65 |
| TempGPol_s (R) | 74 | <-> | Putam (R) | 84 | -77.86 |
| HeschlG (R) | 74 | <-> | Putam (R) | 80 | -74.36 |
| Cing_p (L) | 34 | <-> | Cing_m (R) | 35 | -67.60 |
| Cing_m (R) | 35 | <-> | Cing_p (L) | 34 | -67.60 |
| Olf (L) | 6 | <-> | OrbF_s (R) | 21 | -65.92 |
| Cing_a (R) | 6 | <-> | OrbF_s (R) | 32 | -64.35 |
| PaCLob (L) | 1 | <-> | PreCG (L) | 69 | -58.63 |
| Cing_a (L) | 6 | <-> | OrbF_s (R) | 31 | -58.11 |
| FG_s (L) | 1 | <-> | PreCG (L) | 3 | -57.25 |
| TempG_m (L) | 37 | <-> | Hipp (L) | 85 | -48.71 |
| Thalam (L) | 11 | <-> | OpeFG_i (L) | 77 | -47.34 |
| Caud (L) | 6 | <-> | OrbF_s (R) | 71 | -38.73 |
| Insula (R) | 22 | <-> | Olf (R) | 30 | -38.08 |
| HeschlG (R) | 52 | <-> | Occ_m (R) | 80 | -36.55 |
| Pallid (R) | 28 | <-> | Rectus (R) | 76 | -33.81 |
| FG_s_m (R) | 8 | <-> | FG_m (R) | 24 | -26.14 |
| Insula (L) | 25 | <-> | OrbFG_m (L) | 29 | -25.07 |
| FG_s_m (R) | 6 | <-> | OrbF_s (R) | 24 | -24.98 |
| TempG_m (R) | 48 | <-> | LingG (R) | 86 | -22.83 |
| Cing_a (R) | 14 | <-> | TriFG_i (R) | 32 | -22.72 |
| FG_m (R) | 3 | <-> | FG_s (L) | 8 | -22.63 |
| TempG_s (R) | 52 | <-> | Occ_m (R) | 82 | -22.02 |
| Caud (R) | 2 | <-> | PreCG (R) | 72 | -22.00 |
| Caud (R) | 7 | <-> | FG_m (L) | 72 | -21.47 |
| Thalam (L) | 29 | <-> | Insula (L) | 77 | -20.05 |
| TempGPol_s (R) | 80 | <-> | HeschlG (R) | 84 | -17.98 |
| Caud (L) | 11 | <-> | OpeFG_i (L) | 71 | -12.95 |
| Cing_m (R) | 14 | <-> | TriFG_i (R) | 34 | -11.26 |

| | | | | | |
|---|---|---|---|---|---|
| Cing_m (R) | 1 | <-> | PreCG (L) | 34 | -9.94 |
| RolOpe (L) | 3 | <-> | FG_s (L) | 17 | -9.79 |
| Thalam (L) | 34 | <-> | Cing_m (R) | 77 | -9.29 |
| TempG_i (L) | 81 | <-> | TempG_s (L) | 89 | 12.83 |
| Occ_s (R) | 43 | <-> | Calcar (L) | 50 | 14.42 |
| Thalam (L) | 33 | <-> | Cing_m (L) | 77 | 17.51 |
| TempG_m (L) | 29 | <-> | Insula (L) | 85 | 18.62 |
| Hipp (R) | 4 | <-> | FG_s (R) | 38 | 21.31 |
| SuMargG (R) | 30 | <-> | Insula (R) | 64 | 24.93 |
| Hipp (R) | 10 | <-> | OrbFG_m (R) | 38 | 26.24 |
| PaHippG (L) | 27 | <-> | Rectus (L) | 39 | 29.38 |
| Precun (L) | 39 | <-> | PaHippG (L) | 67 | 36.22 |
| Cing_a (L) | 19 | <-> | SuppMoA (L) | 31 | 36.54 |
| TempG_m (R) | 84 | <-> | TempGPol_s (R) | 86 | 39.53 |
| Calcar (R) | 30 | <-> | Insula (R) | 44 | 41.23 |
| Cing_a (L) | 3 | <-> | FG_s (L) | 31 | 41.93 |
| TempG_i (R) | 84 | <-> | TempGPol_s (R) | 90 | 45.74 |
| Precun (R) | 40 | <-> | PaHippG (R) | 68 | 51.65 |
| Hipp (R) | 22 | <-> | Olf (R) | 38 | 52.88 |
| Caud (L) | 41 | <-> | Amyg (L) | 71 | 53.27 |
| PaHippG (R) | 6 | <-> | OrbF_s (R) | 40 | 59.97 |
| PaHippG (R) | 16 | <-> | OrbFG_i (R) | 40 | 75.49 |
| Amyg (R) | 16 | <-> | OrbFG_i (R) | 42 | 76.48 |
| Calcar (R) | 43 | <-> | Calcar (L) | 44 | 89.78 |
| Calcar (L) | 44 | <-> | Calcar (R) | 43 | 89.78 |
| TempG_m (L) | 81 | <-> | TempG_s (L) | 85 | 91.46 |
| TempG_m (L) | 83 | <-> | TempGPol_s (L) | 85 | 98.73 |
| TempG_i (R) | 86 | <-> | TempG_m (R) | 90 | 103.68 |
| TempG_i (R) | 88 | <-> | TempGPol_m (R) | 90 | 119.24 |
| Pallid (R) | 4 | <-> | FG_s (R) | 76 | 127.12 |
| FusifG (R) | 38 | <-> | Hipp (R) | 56 | 129.35 |
| Cing_a (L) | 23 | <-> | FG_s_m (L) | 31 | 129.51 |
| Amyg (L) | 21 | <-> | Olf (L) | 41 | 161.91 |
| PaHippG (R) | 38 | <-> | Hipp (R) | 40 | 280.90 |
| Pallid (R) | 74 | <-> | Putam (R) | 76 | 325.58 |

**Supp. Table 2. Hubs of the brain for the PBD and HC groups, according to three different classification methods.**

| Hubs | | | | | |
|---|---|---|---|---|---|
| Efficiency | | Connector Hubs | | Provincial Hubs | |
| **PBD** | HC | PBD | HC | PBD | HC |
| L Rolandic Oper | L Rolandic Oper | L Hippocampus | L Hippocampus | L Caudate | L Caudate |
| R Rolandic Oper | R Rolandic Oper | L Precuneus | **R Hippocampus** | R Caudate | R Caudate |
| L Occipital Sup | **R Front Med Orb** | L Putamen | L Precuneus | L Pallidum | L Pallidum |
| L Fusiform | **L Cuneus** | R Cingulum Post | L Putamen | R Pallidum | L Pallidum |
| R Fusiform | L Occipital Sup | **R Occipital Sup** | **R Putamen** | L Cingulum Post | L Cingulum Post |
| L Parietal Inf | **L Occipital Inf** | | R Cingulum Post | L ParaHippocamp | L ParaHippocamp |
| R Parietal Inf | L Fusiform | | **R Amygdala** | R ParaHippocamp | R ParaHippocamp |
| R Heschl | R Fusiform | | | L Fusiform | L Fusiform |
| **R Temp Pole m** | L Parietal Inf | | | R Fusiform | R Fusiform |
| | R Parietal Inf | | | R Parietal Inf | R Parietal Inf |
| | R Heschl | | | **R Putamen** | **L Frontal Sup Orb** |
| | **L Temp Inf** | | | **R Insula** | **R Frontal Sup Orb** |
| | **R Temp Inf** | | | | **L Postcentral** |
| | | | | | **R Heschl** |

**Supp. Table 3. Partial Correlation coefficients between nodes with significant $E_{nodal}$ group-difference and different neurocognitive/mood scores, for both populations.**

| Network metric | Group | Partial Correlation Coefficient | | | | | |
|---|---|---|---|---|---|---|---|
| | | VIQ | PIQ | FSIQ | Coding | Positive | Negative |
| $E_{nodal}$ of ' L Front Sup Orb ' | HC | 0.07 | 0.01 | 0.06 | - | - | - |
| | PBD | 0.38 | 0.23 | 0.36 | 0.41 | 0.13 | -0.31 |
| $E_{nodal}$ of ' L Front Mid Orb ' | HC | 0.11 | -0.17 | 0.03 | - | - | - |
| | PBD | 0.02 | -0.35 | -0.21 | -0.21 | 0.24 | 0.57 * |
| $E_{nodal}$ of ' L Front Inf Orb ' | HC | 0.30 | 0.31 | 0.36 | - | - | - |
| | PBD | 0.74 * | 0.60 * | 0.77 * | 0.15 | -0.34 | -0.50 |
| $E_{nodal}$ of ' R Rolandic Oper ' | HC | 0.12 | 0.22 | 0.18 | - | - | - |
| | PBD | 0.18 | 0.30 | 0.27 | 0.52 | 0.39 | 0.07 |
| $E_{nodal}$ of ' R Front Med Orb ' | HC | 0.35 | 0.43 | 0.42 | - | - | - |
| | PBD | 0.21 | 0.04 | 0.14 | 0.04 | -0.33 | -0.09 |
| $E_{nodal}$ of ' R Insula ' | HC | -0.09 | -0.03 | -0.07 | - | - | - |
| | PBD | 0.18 | 0.34 | 0.31 | 0.40 | 0.19 | -0.09 |
| $E_{nodal}$ of ' L Occipital Inf ' | HC | -0.07 | 0.27 | 0.07 | - | - | - |
| | PBD | -0.25 | 0.28 | 0.00 | 0.16 | -0.12 | 0.19 |
| $E_{nodal}$ of ' R SupraMarginal ' | HC | 0.08 | 0.33 | 0.20 | - | - | - |
| | PBD | -0.05 | -0.04 | -0.11 | 0.11 | -0.01 | -0.04 |
| $E_{nodal}$ of ' R Temporal Inf ' | HC | -0.11 | 0.00 | -0.08 | - | - | - |
| | PBD | 0.33 | 0.34 | 0.40 | 0.17 | -0.34 | -0.26 |
| $E_{nodal}$ of ' R Cingulum Mid ' | HC | 0.36 | -0.04 | 0.25 | - | - | - |
| | PBD | 0.23 | 0.46 | 0.36 | 0.45 | -0.14 | -0.24 |
| $E_{nodal}$ of ' R Heschl ' | HC | 0.24 | 0.22 | 0.26 | - | - | - |
| | PBD | -0.55 | -0.32 | -0.53 | -0.12 | 0.28 | 0.20 |

**Supp. Table 4. Partial Correlation coefficients between global graph theory metrics and neurocognitive/mood scores, for both populations.**

| | | Partial Correlation Coefficient | | | | | |
|---|---|---|---|---|---|---|---|
| Network metric | Group | VIQ | PIQ | FSIQ | Coding | Positive | Negative |
| Mean Connectivity *[1] | HC | 0,067 | -0,010 | 0,037 | - | - | - |
| | PBD | 0,372 | 0,148 | 0,286 | 0,316 | 0,202 | -0,320 |
| Total Number of Fibers | HC | 0,121 | 0,439 | 0,288 | - | - | - |
| | PBD | 0,370 | 0,338 | 0,395 | 0,341 | -0,216 | -0,172 |
| Fibers per Connection | HC | 0,349 | 0,274 | 0,373 | - | - | - |
| | PBD | 0,282 | 0,467 | 0,398 | 0,256 | 0,053 | -0,249 |
| Degree | HC | -0,101 | 0,260 | 0,050 | - | - | - |
| | PBD | 0,310 | 0,173 | 0,280 | 0,300 | -0,296 | -0,086 |
| Connection density | HC | -0,101 | 0,260 | 0,050 | - | - | - |
| | PBD | 0,310 | 0,173 | 0,280 | 0,300 | -0,296 | -0,086 |
| Average Clustering *[2] | HC | -0,072 | -0,179 | -0,132 | - | - | - |
| | PBD | -0,175 | -0,365 | -0,289 | -0,442 | 0,179 | 0,019 |
| Characteristic Path Length *[2] | HC | -0,230 | 0,161 | -0,077 | - | - | - |
| | PBD | 0,159 | 0,573 | 0,369 | 0,034 | -0,056 | -0,536 |
| Small World | HC | 0,004 | -0,221 | -0,102 | - | - | - |
| | PBD | -0,213 | -0,512 | -0,382 | -0,467 | 0,200 | 0,143 |
| Global Efficiency *[1] | HC | 0,032 | 0,094 | 0,062 | - | - | - |
| | PBD | 0,347 | 0,146 | 0,283 | 0,393 | -0,116 | -0,187 |
| Local Efficiency *[1] | HC | -0,011 | -0,056 | -0,038 | - | - | - |
| | PBD | 0,401 | 0,216 | 0,337 | 0,337 | 0,248 | -0,250 |
| Modularity | HC | 0,055 | 0,299 | 0,168 | - | - | - |
| | PBD | -0,229 | -0,099 | -0,242 | -0,139 | 0,445 | -0,183 |
| Modules | HC | -0,205 | -0,327 | -0,285 | - | - | - |
| | PBD | -0,138 | -0,279 | -0,276 | 0,079 | 0,014 | -0,193 |

*[1] Calculated on the normalised version of the SC matrices ([0,1]).

*[2] Weigted version; divided by random networks; uses the original SC matrices.

**Supp. Table 5. Group differences in graph theoretical metrics.**

| Network metric | Controls | | Bipolars | | MW U-test | KS-test | t-test |
|---|---|---|---|---|---|---|---|
| | mean | SD | mean | SD | pvalue | | |
| Mean Connectivity *[1] | 0,119 | 0,005 | 0,120 | 0,007 | 0,619 | 0,589 | 0,818 |
| Total Number of Fibers | 422208,090 | 32796,155 | 412160,037 | 47764,226 | 0,534 | 0,890 | 0,507 |
| Fibers per Connection | 328,996 | 14,624 | 326,791 | 18,328 | 0,678 | 0,890 | 0,718 |
| Degree | 1284,667 | 99,853 | 1260,400 | 121,498 | 0,820 | 0,998 | 0,555 |
| Connection density | 0,160 | 0,012 | 0,157 | 0,015 | 0,820 | 0,998 | 0,555 |
| Average Clustering *[2] | 3,742 | 0,332 | 3,803 | 0,480 | 0,901 | 0,998 | 0,691 |
| Characteristic Path Length *[2] | 1,559 | 0,041 | 1,529 | 0,042 | 0,062 | 0,136 | 0,054 |
| Small World | 2,401 | 0,206 | 2,487 | 0,296 | 0,534 | 0,890 | 0,361 |
| Global Efficiency *[1] | 0,075 | 0,005 | 0,075 | 0,007 | 0,967 | 0,998 | 0,930 |
| Local Efficiency *[1] | 0,092 | 0,004 | 0,093 | 0,005 | 0,678 | 0,589 | 0,820 |
| Modularity | 0,596 | 0,012 | 0,594 | 0,016 | 0,740 | 0,589 | 0,735 |
| Modules | 7,133 | 0,516 | 7,267 | 0,458 | - | - | - |

*[1] Calculated using the normalised version of the SC matrices ([0,1]).

*[2] Weighted version; divided by random networks; uses the original SC matrices.
