## Supplementary Figure for "Disrupted brain structural connectivity in Pediatric Bipolar Disorder with psychosis"

### Supplementary Figures

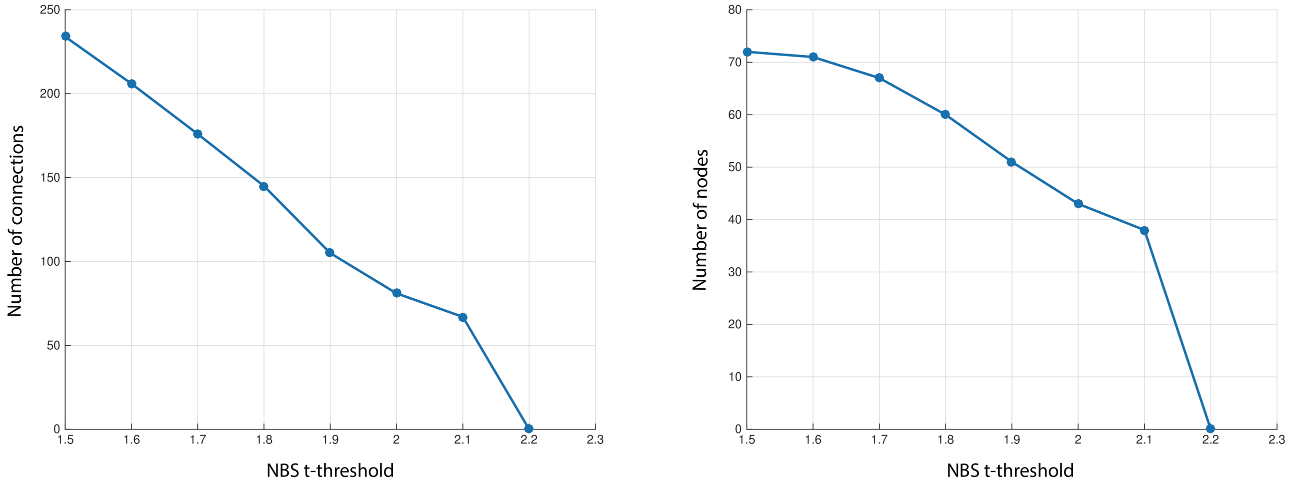


**Supp Figure 1. Network-based statistics.** Relationship between t-threshold and number of connections/nodes, for the largest anatomical component. The t-threshold used in this study (*2.1*) was selected based on the maximal t-threshold ≥ 2.0 where a component was found, and generated an NBS component with approximately 43% nodes of the network and 71links.
